# Evidence that faecal carriage of resistant *Escherichia coli* by 16-week-old dogs in the United Kingdom is associated with raw feeding

**DOI:** 10.1101/2021.04.17.440283

**Authors:** Oliver Mounsey, Kezia Wareham, Ashley Hammond, Jacqueline Findlay, Virginia C. Gould, Katy Morley, Tristan A. Cogan, Katy M.E. Turner, Matthew B. Avison, Kristen K. Reyher

## Abstract

We report a survey (August 2017 to March 2018) and risk factor analysis of faecal carriage of antibacterial-resistant (ABR) *Escherichia coli* in 223 sixteen-week-old dogs in the United Kingdom. Raw feeding was associated with the presence of *E. coli* resistant to fluoroquinolones, tetracycline, amoxicillin, and streptomycin, but not to cefalexin or cefotaxime. Whole genome sequencing of 30 fluoroquinolone-resistant (FQ-R), 22 cefotaxime-resistant (CTX-R) and seven dual FQ-R/CTX-R *E. coli* isolates showed a wide range of sequence types (STs), an approximately 50:50 split of CTX-M:AmpC-mediated CTX-R, and almost exclusively mutational FQ-R dominated by ST744 and ST162. Comparisons between *E. coli* isolates from puppies known to be located within a 50 x 50 km region with those isolated from human urinary tract and bloodstream infections (isolated in parallel in the same region) identified a clone of ST963 *E. coli* carrying chromosomal *bla*_CMY.2_ in two puppies and causing two urinary tract infections and one bloodstream infection. Furthermore, an ST744 FQ-R clone was carried by one puppy and caused one urinary tract infection. Accordingly, we conclude that raw feeding is associated with carriage of ABR *E. coli* in dogs even at sixteen weeks of age and that bacteria carried by dogs are shared with humans.

## Introduction

Antimicrobial resistance and particularly antibacterial resistance (ABR) has many negative impacts on the health and welfare of humans and animals including increased morbidity and mortality and an increase in treatment costs (1). ABR is linked across human populations, animal populations and the environment, and it is possible for ABR bacteria – or ABR genes that they carry – to be passed between these realms (2). Previous research has indicated that farmed animals act as reservoirs of ABR bacteria that can be transmitted to humans either through the food chain, through direct contact between humans and animals or via the environment (3,4).

In many countries, particularly in urban areas, interaction between humans and farm animals – directly or via the environment – is limited. This may explain why studies using whole genome sequencing (WGS) have found little evidence that sharing of ABR bacteria between farmed animals and humans is a significant problem (5–8). However, close interaction between humans and domestic animals is common in such areas. Accordingly, it may be that for many people around the world, a pet dog is a more likely source of ABR bacteria than are farmed animals. Indeed, ABR bacteria found in domestic pets and their owners are often indistinguishable (9–12). A key ABR pathogen of relevance is *Escherichia coli,* which is carried in the intestines of humans, farmed and companion animals, and causes a significant disease burden in all three, and especially in humans (13).

There are several ways that dogs may become colonised by ABR *E. coli* and so bring them into the home. Ingestion is an essential part of colonisation; therefore, ingestion of faeces or faecally contaminated food or water by dogs may be a key source of ABR bacteria derived from humans and farmed animals. For example, farm animal manure is often spread on pastureland where dogs might be exercised. Wastewater from farm run-off or from human sewage outlets may introduce *E. coli* to fresh and sea water where dogs might bathe (14,15). Meat can be contaminated with animal faeces during slaughter, and if eaten in its raw form by a dog, may lead to *E. coli* colonisation (16). Research has also suggested that dogs become colonised by ABR bacteria when visiting veterinary hospitals, which act as reservoirs for multidrug resistant (MDR) organisms, and particularly if the dog receives antibacterial therapy (17–19). Recent research examining 374 veterinary practices in the UK estimated that during the two years investigated, around 25% of approximately one million pet dogs registered received at least one antibacterial course. Of dog antibacterial usage in this study, 60% was classified as use of a ‘critically important’ medicine as defined by WHO criteria (20).

Overall, ABR bacteria have been detected in both healthy and sick adult dogs and associations have been found between increased carriage of ABR bacteria and exposure to antibacterials (19). Associations have also been found between increased carriage of ABR bacteria following veterinary healthcare in general as well as with coprophagia and with the feeding of raw poultry (21–25). Of direct relevance to the present study, two UK studies have identified associations between ABR in faecal *E. coli* of adult dogs and those dogs being fed raw meat (21,24).

Up to now, there has not been any published work reporting very early life risk factors for carriage of ABR *E. coli* in domestic pet dogs. In the UK, current recommendations are for juvenile dogs to be weaned onto solid food and receive a core vaccination at six to eight weeks of age and then receive booster vaccinations every two to four weeks until 16 weeks of age (26). Dogs should stay with their mother until eight weeks of age, and owners are usually advised not to walk their dog outside in public places until after the dog has had its second vaccination (approximately 12 weeks of age).

In this study, risk factors were investigated to explore associations between various lifestyle factors and the detection of ABR *E. coli* in faecal samples taken from dogs at 16 weeks of age. Practices and behaviours that might increase ingestion of faecal bacteria from the environment or food were particularly considered. Furthermore, WGS was used to characterise ABR isolates. The focus was specifically on resistance to critically important antibacterials: 3^rd^ generation cephalosporins (3GC), e.g., cefotaxime (CTX) and fluoroquinolones. CTX resistant (CTX-R) and fluoroquinolone resistant (FQ-R) *E. coli* carried by a sub-set of puppies were compared with those cultured from human urinary tract and bloodstream infections collected in parallel within the same 50 x 50 km region, to investigate whether there is evidence of transmission.

## Results and Discussion

### *Risk factors for carriage of ABR* E. coli *in dogs at 16 weeks of age*

In total, 295 dogs were recruited and data for 223 dogs were included in the analysis. Submissions were excluded if the questionnaire was not fully completed (n=14) or because the faecal sample did not grow enough *E. coli* to be sure of ABR status as defined in Experimental (n=58). For each of the 223 included faecal samples, ABR *E. coli* carriage status was categorised as positive or negative for resistance to five test antibacterials: amoxicillin, cefalexin, ciprofloxacin, streptomycin, or tetracycline, as set out in Experimental. In a preliminary Chi-squared analysis, the only significant risk factor identified for 16-week-old dogs providing faecal samples carrying *E. coli* resistant to at least one antibacterial was having been fed raw food *(p* <0.001; **Table 1**). Subsequent univariable and multivariable logistic regression analyses showed a strong association between raw feeding and carriage of *E. coli* resistant to any one of the five antibacterials tested as well as individually with resistance to each of the antibacterials tested except cefalexin (**Table 2**).

**Table 1.** Baseline data for all 16-week-old dogs (n=223) and associations with risk factors for carriage of *E. coli* resistant to at least one test antibacterial. *p*-values were calculated using the Pearson Chi-squared test (Stata/IC 15.1, StataCorp LLC, College Station, TX, USA). The bold figures show a *p*-value < 0.05.

| Risk factor from questionnaire | Response to question | Response to question total (n=223) | Also resistant to at least one antibiotic (n=106) | *p*-value |
| --- | --- | --- | --- | --- |
| Fed raw food | Yes | 43 | 32/43 | <0.001 |
|  | No | 180 | 76/180 |  |
| Walked in town | Yes | 181 | 84/181 | 0.21 |
|  | No | 42 | 24/42 |  |
| Walked on farmland | Yes | 142 | 69/142 | 0.95 |
|  | No | 81 | 39/81 |  |
| Walked on beaches | Yes | 103 | 52/103 | 0.57 |
|  | No | 120 | 56/120 |  |
| Walked in the countryside | Yes | 191 | 95/191 | 0.34 |
|  | No | 32 | 13/32 |  |
| Walking near cattle | Yes | 84 | 37/70 | 0.31 |
|  | No | 139 | 71/139 |  |
| Swum/ paddled/ played in salt water | Yes | 62 | 32/62 | 0.56 |
|  | No | 161 | 76/161 |  |
| Swum/ paddled/ played in lake water | Yes | 29 | 17/29 | 0.24 |
|  | No | 194 | 91/194 |  |
| Swum/ paddled/ played in river water | Yes | 66 | 33/66 | 0.76 |
|  | No | 157 | 75/157 |  |
| Swum/ paddled/ played in pond water | Yes | 65 | 38/65 | 0.06 |
|  | No | 158 | 70/158 |  |

**Table 2.** Univariable and multivariable logistic regression analyses using questionnaire data and antibacterial-resistant *E. coli* data for 16-week-old dogs (n=223). Presentation: Odds ratio (95% confidence interval) *p-*value. Only risk actors significantly associated with resistance (*p-*value < 0.05) are included.

| Risk Factor | Univariable<br>(n=223) | Multivariable for all samples<br>(n=223) |
| --- | --- | --- |
| **Resistance to at least one antibacterial (n=108)** |  |  |
| Fed raw food | 3.98 (1.89 to 8.40)<br><0.001 | 3.98 (1.89 to 8.40) <0.001 |
| **Resistance to ciprofloxacin (n=26)** |  |  |
| Fed raw food | 12.42 (5.01 to 30.78) <0.001 | 12.42 (5.01 to 30.78) <0.001 |
| **Resistance to tetracycline (n=81)** |  |  |
| Fed raw food | 4.47 (2.21 to 9.05)<br><0.001 | 4.47 (2.21 to 9.05) <0.001 |
| **Resistance to amoxicillin (n=93)** |  |  |
| Fed raw food | 3.30 (1.64 to 6.63)<br>0.001 | 3.18 (1.57 to 6.42) 0.001 |
| Swam/paddled/<br>played in pond<br>water | 2.01 (1.12 to 3.61)<br>0.02 | 1.91 (1.05 to 3.48) 0.04 |
| **Resistance to cephalexin (n=34)** |  |  |
| No statistically significant risk factors identified |  |  |
| **Resistance to streptomycin (n=51)** |  |  |
| Fed raw food | 8.23 (3.95 to 17.15) <0.001 | 8.23 (3.95 to 17.15) <0.001 |

The most substantial risk associated with raw feeding in 16-week-old dogs was that of carriage of FQ-R *E. coli* (**Table 2**). This association has previously been reported in adult dogs in the UK; a study based on 445 dogs found that feeding raw poultry significantly increased the risk of carrying FQ-R *E. coli* in faeces (22). Findings from the present study extend these earlier studies to show that the impact of raw feeding on ABR *E. coli* carriage can be seen as early as 10 weeks after the first introduction of solid food. Faecal samples taken from broilers at a slaughterhouse commonly contain FQ-R *E. coli* (27) and raw chicken imported into (28) and produced in the UK (29) have been identified as contaminated with FQ-R *E. coli.* Feeding raw chicken could therefore be a source of FQ-R *E. coli* in our study, as has been seen with adult dogs (22), but this remains to be confirmed. The risk of dogs acquiring ABR bacteria from meat would be mitigated simply by cooking that meat to reduce any contamination with faecal bacteria that occurs at slaughter and during processing.

### *Molecular epidemiology of CTX-R and FQ-R* E. coli *from puppies*

Of faecal samples from 34 dogs that contained cefalexin-resistant *E. coli,* 27 gave CTX-R isolates. PCR analysis was used to identify mobile resistance genes associated with CTX-R in these isolates; where the same PCR profile was seen for multiple CTX-R isolates from a sample, a single isolate was taken forward for WGS to represent that CTX-R type and sample. In total, 29 unique isolates from these 27 dogs were analysed by WGS. Of these, seven isolates were also FQ-R (**Table 3**).

**Table 3.** Characterisation of CTX-R *E. coli* from puppies using WGS. Stars denote locally recruited dogs. Bold underlining denotes dogs fed raw food.

| Dog ID | *E. coli* ST | FQ-R mechanism(s) | CTX-R mechanism |
| --- | --- | --- | --- |
| DOG 1 | ST372 |  | CMY-2 |
| DOG 2 | ST10 |  | CTX-M-1 |
| DOG 3** | ST2179 | *gyrA* S83L; *parC* S80I | CTX-M-65 |
| DOG 4 | ST744 | *gyrA* S83L; *gyrA* D87N; *parC* A56T; *parC* S80I | CTX-M-1 |
| DOG 5 | ST38 | (*gyrA* S83L; *aac*(6')-Ib-cr) | CTX-M-15 |
| DOG 6 | ST58 |  | *ampC* -42C>T |
| DOG 7 | ST88 |  | CTX-M-1 |
| **<u>DOG 8</u>** | ST88 |  | *ampC* -42C>T |
| DOG 9 | ST38 | (*qnrS1*) | CTX-M-15 |
| DOG 10 | ST963 |  | CMY-2 |
| DOG 11 | ST1196 | *gyrA* S83L; *gyrA* D87N; *parC* S80I; *qnrB4* | DHA-1 |
| DOG 12 | ST215 | (*qnrS1*) | CTX-M-15 |
| DOG 13 | ST973 |  | CMY-2 |
| DOG 15 | ST6096 |  | CMY-2 |
| DOG 16 | ST3889 | (*qnrS1*) | CTX-M-15 |
| **<u>DOG 18</u>** | ST744 | *gyrA* S83L; *gyrA* D87N; *parC* A56T; *parC* S80I | CTX-M-1 |
| DOG 21** | ST69 | (*qnrS1*) | CTX-M-14 |
| DOG 21** | ST963 |  | CMY-2 |
| DOG 22** | ST963 |  | CMY-2 |
| DOG 23 | ST744 | *gyrA* S83L; *gyrA* D87N; *parC* A56T; *parC* S80I | CTX-M-1 |
| DOG 25 | ST155 |  | *ampC* -42C>T |
| **<u>DOG 27</u>** | ST88 | (*gyrA* S83L) | *ampC* -42C>T |
| **<u>DOG 27</u>** | ST1431 | *gyrA* S83L; *gyrA* D87N; *parC* S80I; *qnrS1* | *ampC* -42C>T |
| **<u>DOG 28</u>** | ST602 |  | *ampC* -42C>T |
| DOG 29 | ST4988 | *gyrA* S83L; *gyrA* D87N; *parC* S80I | CTX-M-15 |
| **<u>DOG 31\*\*</u>** | ST88 |  | *ampC* -42C>T |
| DOG 42 | ST1056 |  | CTX-M-1 |
| DOG 43 | ST75 |  | *ampC* -42C>T |
| DOG 44 | ST961 |  | CTX-M-1 |

WGS revealed a wide range of *E. coli* STs and CTX-R mechanisms (**Table 3**): ST88 (one isolate with CTX-M-1; three isolates with mutations in the *ampC* promoter known to be associated with hyper-expression) was dominant, followed by ST744 (three FQ-R isolates with CTX-M-1), ST963 (three isolates with CMY-2) and ST38 (two isolates with CTX-M-15). Seventeen additional isolates, each representing a unique ST, were found to be carrying CTX-M-1 (three isolates), CTX-M-15 (three isolates), CTX-M-65 (one isolate), CTX-M-14 (one isolate), CMY-2 (three isolates) DHA-1 (one isolate) and *ampC* promoter mutation (five isolates).

Overall, therefore, AmpC-type β-lactamase-mediated resistance was found in 15/29 isolates and CTX-M was found in 14/29. This approximately 50:50 split was also seen in a recent analysis of CTX-R *E. coli* from 53 dairy farms in South West England, where amoxicillin/clavulanate use was associated with finding AmpC-mediated CTX-R *E. coli* in farm samples (8). A study examining prescribing at small animal veterinary practices in the UK found that amoxicillin/clavulanate was the most common antibacterial prescribed, accounting for 36% of prescriptions (30), and it has been demonstrated that routine amoxicillin/clavulanate treatment selects for increased CTX-R *E. coli* in the faeces of dogs (19). It could therefore be hypothesised that the reason why clavulanic acid-insensitive AmpC-type β-lactamases are so common in CTX-R *E. coli* carried by dogs is because of high levels of amoxicillin/clavulanate usage in the canine population generally. However, whilst this study did not record veterinary treatments, it seems unlikely that antibacterial therapy was widespread in these puppies, given their age and exclusion of puppies that had been hospitalised. This finding of AmpC dominance is therefore suggestive of transmission into the juvenile dogs in the study. There was no positive association between raw feeding and the presence of CTX-R isolates in general; only six out of 29 CTX-R isolates were from raw-fed dogs (**Table 3**). However, among these, five out of eight of the AmpC hyper-producing isolates were from raw-fed dogs. Whilst these numbers are too small for clear conclusions to be drawn, it is plausible that raw feeding may selectively seed *ampC* hyper-producer *E. coli* carriage.

Carriage of FQ-R *E. coli* was strongly associated with raw feeding in puppies (**Table 2**). From 26 puppies that produced samples carrying FQ-R *E. coli,* 30 isolates were subjected to WGS (**Table 4**) in addition to the seven dual FQ-R/CTX-R isolates discussed above (**Table 3**). Plasmid-mediated quinolone resistance mechanisms (PMQR) were found in only 3/37 FQ-R isolates, and in only one ST58 isolate carrying *qnrS1* and a single *gyrA* mutation (**Table 4**) was there any suggestion that a PMQR was necessary for conferring FQ-R. The other two PMQR-carrying FQ-R isolates were also CTX-R (**Table 3**). These two were an ST1196 isolate carrying *qnrS1* and an ST1431 isolate carrying *qnrB4,* but in both there were also two mutations in *gyrA* and one in *parC,* sufficient to confer FQ-R in the absence of a PMQR gene (31). Indeed, many of the FQ-R isolates collected in this study carried identical mutations and no PMQR genes (**Table 4**). Interestingly, five of the CTX-R isolates that were not FQ-R also carried PMQRs: four had a *qnrS1* gene and one ST38 isolate had an *aac(6)-Ib-cr* gene (**Table 3**). This would support previous conclusions that carriage of these genes is not sufficient to confer FQ-R in the absence of other mechanisms (31).

**Table 4.** Characterisation of FQ-R *E. coli* from puppies using WGS. Stars denote locally recruited dogs. Bold underlining denotes dogs fed raw food.

| Dog ID | *E. coli* ST | FQ-R mechanism(s) |
| --- | --- | --- |
| DOG 4 | ST744 | *gyrA* S83L; *gyrA* D87N; *parC* A56T; *parC* S80I |
| **<u>DOG 9</u>** | ST162 | *gyrA* S83L; *gyrA* D87N; *parC* S80I |
| DOG 10 | ST224 | *gyrA* S83L; *gyrA* D87N; *parC* S80I |
| DOG 14 | ST162 | *gyrA* S83L; *gyrA* D87N; *parC* S80I |
| **<u>DOG 16</u>** | ST744 | *gyrA* S83L; *gyrA* D87N; *parC* A56T; *parC* S80I |
| **<u>DOG 17</u>** | ST453 | *gyrA* S83L; *gyrA* D87N; *parC* S80I |
| **<u>DOG 17</u>** | ST58 | *gyrA* S83L; *qnrS1* |
| **<u>DOG 17</u>** | ST744 | *gyrA* S83L; *gyrA* D87N; *parC* A56T; *parC* S80I |
| **<u>DOG 18</u>** | ST224 | *gyrA* S83L; *gyrA* D87N; *parC* S80I |
| **<u>DOG 19</u>** | ST744 | *gyrA* S83L; *gyrA* D87N; *parC* A56T; *parC* S80I |
| **<u>DOG 20</u>** | ST162 | *gyrA* S83L; *gyrA* D87N; *parC* S80I |
| DOG 21** | ST10 | *gyrA* S83L; *gyrA* D87N; *parC* S80I |
| **<u>DOG 24</u>** | ST744 | *gyrA* S83L; *gyrA* D87N; *parC* A56T; *parC* S80I |
| **<u>DOG 26</u>** | ST1196 | *gyrA* S83L; *gyrA* D87N; *parC* S80I |
| **<u>DOG 26</u>** | ST1011 | *gyrA* S83L; *gyrA* D87N; *parC* S80I |
| **<u>DOG 30</u>** | ST744 | *gyrA* S83L; *gyrA* D87N; *parC* A56T; *parC* S80I |
| **<u>DOG 31\*\*</u>** | ST744 | *gyrA* S83L; *gyrA* D87N; *parC* A56T; *parC* S80I |
| DOG 32** | ST162 | *gyrA* S83L; *gyrA* D87N; *parC* S80I |
| DOG 33 | ST542 | *gyrA* S83L; *parC* S80I |
| **<u>DOG 34</u>** | ST1011 | *gyrA* S83L; *gyrA* D87N; *parC* S80I |
| **<u>DOG 34</u>** | ST6817 | *gyrA* S83L; *gyrA* D87N; *parC* S80I |
| DOG 35 | ST1011 | *gyrA* S83L; *gyrA* D87N; *parC* S80I |
| DOG 35 | ST224 | *gyrA* S83L; *gyrA* D87N; *parC* S80I |
| **<u>DOG 36</u>** | ST155 | *gyrA* S83L; *gyrA* D87N; *parC* S80I |
| DOG 37 | ST744 | *gyrA* S83L; *gyrA* D87N; *parC* A56T; *parC* S80I |
| DOG 37 | ST162 | *gyrA* S83L; *gyrA* D87N; *parC* S80I |
| DOG 38** | ST744 | *gyrA* S83L; *gyrA* D87N; *parC* A56T; *parC* S80I |
| DOG 39** | ST162 | *gyrA* S83L; *gyrA* D87N; *parC* S80I |
| DOG 40 | ST1193 | *gyrA* S83L; *gyrA* D87N; *parC* S80I; *parE* L416F |
| DOG 41** | ST1011 | *gyrA* S83L; *gyrA* D87N; *parC* S80I |

Of the FQ-R isolates sequenced, ST744 (12/37 isolates) dominated, with 6/37 isolates identified as ST162, 4/37 identified as ST1011, 3/37 identified as ST224, 2/37 identified as ST1196 and individual examples of 10 other STs (**Table 3, 4**).

### *Evidence of faecal carriage of CTX-R and FQ-R* E. coli *in puppies that are clonally related to those causing urinary and bloodstream infections in humans in the same geographical area*

A phylogenetic analysis of all the CTX-R and FQ-R isolates from puppies subjected to WGS in this study was constructed (**Figure 1**). There were three clusters of isolates with chromosomal mutations conferring resistance: FQ-R isolates of ST162 and ST744 with multiple gyrase and topoisomerase mutations and a smaller ST88 cluster with chromosomal *ampC* promoter mutations conferring CTX-R and amoxicillin/clavulanate resistance. In contrast, mobile resistance mechanisms were spread widely across the phylogenetic tree. Notably, one FQ-R isolate was ST1193, which is an important clone currently emerging in human infections and of the most pathogenic phylogroup, B2 (32). It was therefore interesting to test relationships between CTX-R and FQ-R isolates from locally recruited dogs with human urinary CTX-R and FQ-R isolates from people living in the same geographical area as the locally recruited dogs (33,34) whose infections occurred within the same six-month period as collection of the canine faecal samples yielding these isolates.

**Figure 1:**
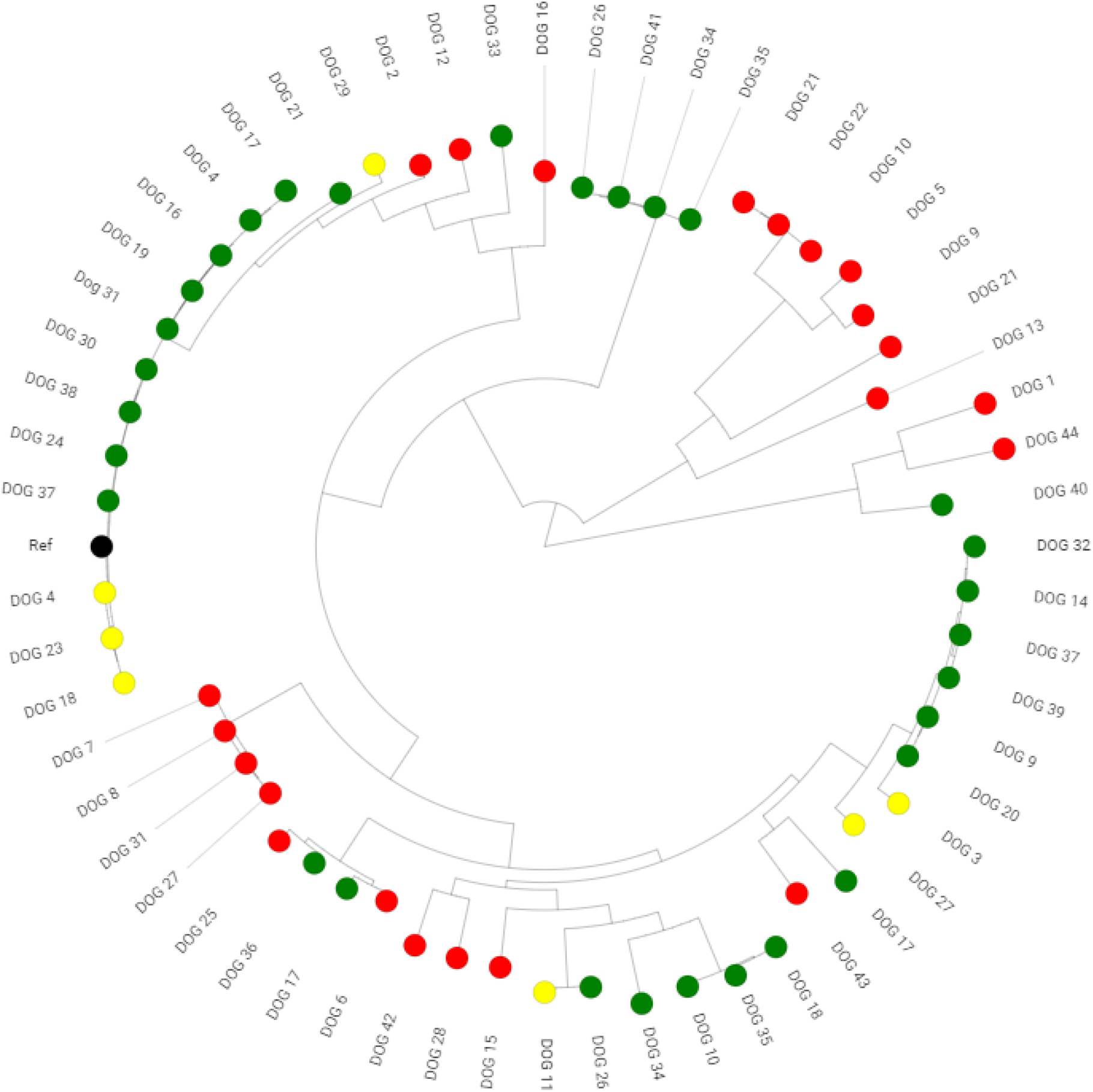
Core genome phylogenetic analysis of antibacterial-resistant *E. coli* from puppies. CTX-R isolates are labelled red, FQ-R isolates are labelled green and CTX-R/FQ-R dual-resistant isolates are labelled yellow. The randomly assigned Dog ID relevant to each isolate is also labelled.

There were four CTX-R isolates from locally recruited dogs; two of these (from two different dogs: Dog 21 and Dog 22) were ST963; the others were ST88 and ST2179 (**Table 3**). None of the 225 CTX-R urinary *E. coli* (33) in the comparison was ST2179 and a SNP distance analysis showed that the canine ST88 isolate was >1000 SNPs distant in the core genome from its closest ST88 human urinary isolate. A core genome SNP distance of 30 or fewer is commonly seen in Enterobacteriales isolates that are confirmed to be part of an acute outbreak of foodborne illness (35). Hence, for these ST88 isolates, there was no evidence for sharing of isolates between dogs and humans. In contrast, the two canine ST963 isolates were 37 SNPs different from each other, suggesting recent sharing of the isolate. Significantly, however, the isolate from Dog 21 was <50 SNPs different from each of two human urinary ST963 isolates, and the isolate from Dog 22 was <65 SNPs from these same two human urinary isolates. Even more troubling, the isolate from Dog 21 was only 34 SNPs different from a CTX-R ST963 bloodstream isolate, one of 82 CTX-R bloodstream isolates collected in parallel from clinical cases in the same geographical region at the same time. The isolate from Dog 22 was 51 SNPs different from this bloodstream isolate. The urinary and bloodstream isolates were between 31 and 38 SNPs different from each other, so this is clear evidence for sharing of the human and canine CTX-R ST963 isolates. Each of these isolates (two canine, two urinary and one bloodstream) had a mobile *bla*_CMY-2_ gene embedded into the chromosome at the same position – proximal to *nhaRA, dnaJ* – which is further evidence of descent from a recent common ancestor. Most interestingly, another canine ST963 isolate was identified in this study, but not in a locally recruited dog (Dog 10, **Figure 1**). In this case, the isolate was 33, 35 and 21 SNPs different from the two urinary isolates and the bloodstream isolate, respectively, an even closer match than that seen with isolates from the two locally recruited dogs, suggesting even more recent sharing. Whilst Dog 10 was not locally recruited, it is possible that it could still be based locally as address details for the nationally recruited dogs were not available for analysis.

Of the seven FQ-R isolates from locally recruited puppies (**Table 3**, **Table 4**), five were of STs found amongst 188 FQ-R urinary *E. coli* from people living in the same geographical area, isolated within six months of collection of the isolates from puppies (34). Of the canine isolates, one was ST10 and two each were ST744 and ST162 (**Table 4**). One of the ST744 isolates was 47 SNPs different from a human urinary isolate, which is suggestive of sharing, as defined above. Among the other four canine, the lowest SNP difference from a human isolate was 324, which does not suggest sharing in these cases. Interestingly, the puppy carrying the seemingly shared ST744 isolate, Dog 31, was the only FQ-R *E. coli* positive locally recruited dog reported to be fed raw meat (**Table 4**).

## Conclusions

This study has identified raw meat feeding as a risk factor for the excretion of ABR *E. coli* in the faeces of 16-week-old puppies, with particularly strong impact on excretion of isolates resistant to the critically important fluoroquinolones. If owners insist on feeding raw meat to their dog, it is essential that they fully understand this practice puts their dog at risk of becoming colonised with bacteria resistant to critically important antibacterials.

*E. coli* is the most clinically important opportunistic human bacterial pathogen (13). ABR *E. coli* infections are more difficult to treat, and result in more morbidity and higher mortality rates (13); there is also strong evidence that domestic pet dogs transmit ABR bacteria to humans (9–12, 36,37) and this study provides clear evidence of the faecal carriage within puppies of CTX-R and FQ-R *E. coli* clonally related to those that have also caused urinary and bloodstream infections in humans living in the same geographical region collected within months of each other. Therefore, if owners feed raw food to their dog, practices that mitigate the risk of onward transmission of ABR *E. coli* – which are more likely to be carried by these dogs – to humans should be encouraged. These include strict hygiene practices when anyone (particularly those vulnerable to bacterial infection) interacts with a raw-fed dog along with scrupulous disposal of the dog’s faeces so that it cannot pose a risk to the general human population by contaminating the wider environment with ABR *E. coli.*

## Experimental

### Recruitment of the cohorts

Dog owners were recruited to take part in this study in two ways: (i) 236 were already recruited to the Dogs Trust “Generation Pup” project, a longitudinal study examining the health, welfare and behaviour of dogs across the UK (39) and (ii) 59 were locally recruited via word-of-mouth advertisement to clients bringing young dogs in for routine checks to veterinary practices in Somerset and Bristol, via puppy socialisation classes and via social media as well as local media advertisement. Locally recruited owners answered survey questions (listed in **Table 1**). As part of Generation Pup, owners completed more extensive surveys relating to their dogs at 16 weeks of age and responses to relevant survey questions (**Table 1**) were extracted from wider Generation Pup survey data. All dog owners also supplied a single faecal sample collected from their dog at 16 weeks of age. All dog owners were recruited between August 2017 and March 2018, and all owners gave consent. Ethical approval for this study was granted by the University of Bristol Health Sciences Student Research Ethics Committee (56783). Health status of the dogs and prior veterinary treatment was not recorded for locally recruited dogs, and so was not included in the analysis. However, dogs that had been previously hospitalised were excluded.

### Faecal samples and processing

All dog owners were supplied with a sample collection pack comprised of a specimen bottle, gloves, biohazard bag and a freepost envelope. Faecal samples were sent by post to the University of Bristol’s Veterinary School alongside the consent form and, for locally recruited dogs, a questionnaire. To process each faecal sample, approximately 0.1-0.5 g of faeces was taken and weighed. Ten millilitres per gram of phosphate buffered saline (PBS) was added to the sample and the mixture vortexed. Next, 0.5 mL of the faecal/PBS homogenate was added to 0.5 mL of 50% v/v sterile glycerol and processed as below.

### Testing for ABR bacteria

Data were collapsed into a binary “positive/negative” outcome for the homogenate derived from each faecal sample. ABR positivity was defined by the appearance (following 37°C overnight incubation) of blue/green *E. coli* colonies after spreading 20 μL of faecal homogenate (or a 10-fold dilution in PBS if inoculum effect was observed) onto Tryptone Bile X-Glucuronide (TBX) agar plates containing either 0.5 mg/L ciprofloxacin (to identify fluoroquinolone resistance [FQ-R]), 16 mg/L cephalexin, 8 mg/L amoxicillin, 16 mg/L tetracycline, or 64 mg/L streptomycin.

Cefalexin-resistant isolates (up to five per plate) grown from primary processing of faecal samples were sub-cultured onto agar plates containing 2 mg/L of cefotaxime (CTX); isolates that grew were deemed CTX-R and taken forward for further testing. These concentrations were chosen based on relevant human clinical breakpoints as defined by the European Committee on Antimicrobial Susceptibility Testing (40). Faecal homogenates were also plated onto non-antibiotic TBX agar and samples were only included in the study if ≥10 *E. coli* cfu/μL were detected in an undiluted faecal homogenate. Therefore, the limit of detection for ABR for all faecal homogenates included in the analysis was ≤0.5% prevalence.

### Risk factor analysis

Univariable and multivariable logistic regression models were used to evaluate associations between ABR *E. coli* positivity in homogenates derived from faecal samples and risk factors identified from the survey data (Stata/IC 15.1, StataCorp LLC, College Station, TX, USA). A backward stepwise method was used. In this method the full set of possible factors was analysed, with the least significant factors removed one at a time until all remaining factors had *p-*values of 0.05 or less. For the risk factor analysis, questionnaire answers were collapsed into binary ‘Yes/No’ variables; questionnaire answers of ‘sometimes’, ‘often’, ‘almost always’ and ‘frequently’ were all categorised as ‘Yes’.

### Isolates from human infections

WGS data for 225 CTX-R and 188 FQ-R human urinary *E. coli* from a 50 x 50 km region (including the homes of the 59 locally recruited dogs collected during the same timespan as the collection of faecal samples from these puppies) has been reported previously (33, 34). Eighty-two CTX-R *E. coli* bloodstream isolates from patients being treated at hospitals in this same geographical region were obtained from the regional microbiology diagnostic laboratory (Severn Pathology, Southmead Hospital, North Bristol NHS Trust). All infections occurred during the same period as puppy faecal sample collection for this study.

### *PCR and WGS analysis of CTX-R and FQ-R* E. coli

Multiplex PCR assays were used to differentiate CTX-R puppy *E. coli* isolates carrying different β-lactamase genes, as described previously (33). WGS of deduplicated, representative CTX-R and FQ-R isolates from puppies, together with the human CTX-R bloodstream isolates was performed by MicrobesNG (https://microbesng.uk/) on a HiSeq 2500 instrument (Illumina, San Diego, CA, USA) using 2×250 bp paired end reads. Reads were trimmed using Trimmomatic (41) and assembled into contigs using SPAdes (42) 3.13.0 (http://cab.spbu.ru/software/spades/). Contigs were annotated using Prokka (43). ABR genes were assigned using the ResFinder (44) and Sequence Types designated by MLST 2.0 (45) on the Centre for Genomic Epidemiology (http://www.genomicepidemiology.org/) platform. Single nucleotide polymorphism (SNP) distance analysis was performed using SNP-dists (https://github.com/tseemann/snp-dists).

### Phylogenetic analysis

Sequence alignment and phylogenetic analysis was carried out using the Bioconda channel (46) on a server hosted by the Cloud Infrastructure for Microbial Bioinformatics (CLIMB; 47). The reference sequence was *E. coli* ST131 isolate EC958 complete genome (accession: HG941718). Sequences were first aligned to a closed reference sequence and analysed for SNP differences, whilst omitting insertion and deletion elements, using the Snippy alignment program (https://github.com/tseemann/snippy). Alignment was then focused on regions of the genome common to all isolates (the “core genome”) using the Snippy-core program, thus eliminating the complicating factors of insertions and deletions. Aligned sequences were then used to construct a maximum likelihood phylogenetic tree using RAxML utilising the GTRCAT model of rate heterogeneity and the software’s autoMR and rapid bootstrap to find the best-scoring maximum likelihood tree and including tree branch lengths, defined as the number of base substitutions per site compared (48,49). Finally, phylogenetic trees were illustrated using the web-based Microreact program (50).

## Acknowledgements

Genome sequencing was provided by MicrobesNG (http://www.microbesng.uk). The authors thank Dogs Trust and Rachel Casey, Jane Murray, Rachel Kinsman and Michelle Lord for assisting with data collection from the Generation Pup project participants included in this study. The authors are grateful to all puppy owners who took part in this study.

## Funding

This work was funded by grant NE/N01961X/1 to MBA, TAC, KMT, KKR from the Antimicrobial Resistance Cross Council Initiative supported by the seven United Kingdom research councils. Dogs Trust fund the Generation Pup project.

**The authors declare no conflicts of interest.**

## Author Contributions

Conceived the Study: K.K.R., M.B.A.

Collection of Data: K.W. O.M., J.F., K.M. supervised by T.A.C., M.B.A., K.K.R.

Cleaning and Analysis of Data: O.M. K.W. A.H., V.C.G., supervised by M.B.A., K.M.T., K.K.R.

Initial Drafting of Manuscript: K.W., O.M., K. K. R., M.B.A.

Corrected and Approved Manuscript: All Authors

## Notes

### Competing Interest Statement

The authors have declared no competing interest.

## References

1. Friedman ND, Temkin E, Carmeli Y. 2016. The negative impact of antibiotic resistance. Clin Microbiol Infect. 22:416–22.

2. Woolhouse M, Ward M, van Bunnik B, Farrar J. 2015. Antimicrobial resistance in humans, livestock and the wider environment. Philos Trans R Soc Lond B Biol Sci. 370:20140083.

3. Zurfluh K, Glier M, Hächler H, Stephan R. 2015. Replicon typing of plasmids carrying *bla*_CTX_-_M_-_15_ among Enterobacteriaceae isolated at the environment, livestock and human interface. Sci Total Environ. 521-522:75–8.

4. Magouras I, Carmo LP, Stärk KDC, Schüpbach-Regula G. 2017. Antimicrobial Usage and -Resistance in Livestock: Where Should We Focus? Front Vet Sci. 4:148.

5. Findlay J, Mounsey O, Lee WWY, Newbold N, Morley K, Schubert H, Gould VC, Cogan TA, Reyher KK, Avison MB. 2020. Molecular Epidemiology of *Escherichia coli* Producing CTX-M and pAmpCβ-Lactamases from Dairy Farms Identifies a Dominant Plasmid Encoding CTX-M-32 but No Evidence for Transmission to Humans in the Same Geographical Region. Appl Environ Microbiol. 87:e01842–20.

6. Day MJ, Rodríguez I, van Essen-Zandbergen A, Dierikx C, Kadlec K, Schink AK, Wu G, Chattaway MA, DoNascimento V, Wain J, Helmuth R, Guerra B, Schwarz S, Threlfall J, Woodward MJ, Coldham N, Mevius D, Woodford N. 2016. Diversity of STs, plasmids and ESBL genes among *Escherichia coli* from humans, animals and food in Germany, the Netherlands and the UK. J Antimicrob Chemother. 71:1178–1182.

7. Ludden C, Raven KE, Jamrozy D, Gouliouris T, Blane B, Coll F, de Goffau M, Naydenova P, Horner C, Hernandez-Garcia J, Wood P, Hadjirin N, Radakovic M, Brown NM, Holmes M, Parkhill J, Peacock SJ. 2019. One Health Genomic Surveillance of *Escherichia coli* Demonstrates Distinct Lineages and Mobile Genetic Elements in Isolates from Humans versus Livestock. mBio. 10:e02693–18.

8. Alzayn M, Findlay J, Schubert H, Mounsey O, Gould VC, Heesom KJ, Turner KM, Barrett DC, Reyher KK, Avison MB. 2020. Characterization of AmpC-hyperproducing *Escherichia coli* from humans and dairy farms collected in parallel in the same geographical region. J Antimicrob Chemother 75:2471–2479.

9. Toombs-Ruane LJ, Benschop J, French NP, Biggs PJ, Midwinter AC, Marshall JC, Chan M, Drinković D, Fayaz A, Baker MG, Douwes J, Roberts MG, Burgess SA. 2020. Carriage of extended-spectrum beta-lactamase-and AmpC beta-lactamase-producing *Escherichia coli* from humans and pets in the same households. Appl Environ Microbiol. 86:e01613–20.

10. Kidsley AK, White RT, Beatson SA, Saputra S, Schembri MA, Gordon D, Johnson JR, O’Dea M, Mollinger JL, Abraham S, Trott DJ. 2020. Companion Animals Are Spillover Hosts of the Multidrug-Resistant Human Extraintestinal *Escherichia coli* Pandemic Clones ST131 and ST1193. Front Microbiol. 11:1968.

11. Hong JS, Song W, Park HM, Oh JY, Chae JC, Jeong S, Jeong SH. 2020. Molecular Characterization of Fecal Extended-Spectrum β-Lactamase- and AmpCβ-Lactamase-Producing *Escherichia coli* From Healthy Companion Animals and Cohabiting Humans in South Korea. Front Microbiol. 11:674.

12. Hong JS, Song W, Park HM, Oh JY, Chae JC, Shin S, Jeong SH. 2019. Clonal Spread of Extended-Spectrum Cephalosporin-Resistant Enterobacteriaceae Between Companion Animals and Humans in South Korea. Front Microbiol. 10:1371.

13. Adler A, Katz DE, Marchaim D. 2016. The Continuing Plague of Extended-spectrum β-lactamase-producing Enterobacteriaceae Infections. Infect Dis Clin North Am. 30:347–375.

14. Amos GC, Hawkey PM, Gaze WH, Wellington EM. 2014. Waste water effluent contributes to the dissemination of CTX-M-15 in the natural environment. J AntimicrobChemother. 69:1785–91.

15. Leonard AFC, Zhang L, Balfour AJ, Garside R, Hawkey PM, Murray AK, Ukoumunne OC, Gaze WH. 2018. Exposure to and colonisation by antibioticresistant *E. coli* in UK coastal water users: Environmental surveillance, exposure assessment, and epidemiological study (Beach Bum Survey). Environ Int. 114:326–333.

16. Schmidt M, Unterer S, Suchodolski JS, Honneffer JB, Guard BC, Lidbury JA, Steiner JM, Fritz J, Kölle P. 2018. The fecal microbiome and metabolome differs between dogs fed Bones and Raw Food (BARF) diets and dogs fed commercial diets. PLoS One. 13:e0201279.

17. Gibson JS, Morton JM, Cobbold RN, Filippich LJ, Trott DJ. 2011. Risk factors for dogs becoming rectal carriers of multidrug-resistant *Escherichia coli* during hospitalization. Epidemiol Infect. 139:1511–21.

18. Ogeer-Gyles J, Mathews KA, Sears W, Prescott JF, Weese JS, Boerlin P. 2006. Development of antimicrobial drug resistance in rectal *Escherichia coli* isolates from dogs hospitalized in an intensive care unit. J Am Vet Med Assoc. 229:694–9.

19. Schmidt VM, Pinchbeck G, McIntyre KM, Nuttall T, McEwan N, Dawson S, Williams NJ. 2018. Routine antibiotic therapy in dogs increases the detection of antimicrobial-resistant faecal *Escherichia coli*. J Antimicrob Chemother. 73:3305–3316.

20. Prescott JF, Boerlin P. 2016. Antimicrobial use in companion animals and Good Stewardship Practice. Vet Rec. 179:486–488.

21. Wedley AL, Dawson S, Maddox TW, Coyne KP, Pinchbeck GL, Clegg P, Nuttall T, Kirchner M, Williams NJ. 2017. Carriage of antimicrobial resistant *Escherichia coli* in dogs: Prevalence, associated risk factors and molecular characteristics. Vet Microbiol. 199:23–30.

22. Grønvold AM, L’abée-Lund TM, Sørum H, Skancke E, Yannarell AC, Mackie RI. 2010. Changes in fecal microbiota of healthy dogs administered amoxicillin. FEMS Microbiol Ecol. 71:313–26.

23. Leite-Martins LR, Mahú MI, Costa AL, Mendes A, Lopes E, Mendonça DM, Niza-Ribeiro JJ, de Matos AJ, da Costa PM. 2014. Prevalence of antimicrobial resistance in enteric *Escherichia coli* from domestic pets and assessment of associated risk markers using a generalized linear mixed model. Prev Vet Med. 117:28–39.

24. Schmidt VM, Pinchbeck GL, Nuttall T, McEwan N, Dawson S, Williams NJ. 2015. Antimicrobial resistance risk factors and characterisation of faecal *E. coli* isolated from healthy Labrador retrievers in the United Kingdom. Prev Vet Med. 119:31–40.

25. Baede VO, Wagenaar JA, Broens EM, Duim B, Dohmen W, Nijsse R, Timmerman AJ, Hordijk J. 2015. Longitudinal study of extended-spectrum-β-lactamase- and AmpC-producing Enterobacteriaceae in household dogs. Antimicrob Agents Chemother. 59:3117–24.

26. Day MJ, Horzinek MC, Schultz RD, Squires RA; Vaccination Guidelines Group (VGG) of the World Small Animal Veterinary Association (WSAVA). 2016. WSAVA Guidelines for the vaccination of dogs and cats. J Small AnimPract. 57:E1–E45.

27. Costa D, Vinué L, Poeta P, Coelho AC, Matos M, Sáenz Y, Somalo S, Zarazaga M, Rodrigues J, Torres C. 2009. Prevalence of extended-spectrum beta-lactamase-producing *Escherichia coli* isolates in faecal samples of broilers. Vet Microbiol. 138:339–44.

28. Warren RE, Ensor VM, O’Neill P, Butler V, Taylor J, Nye K, Harvey M, Livermore DM, Woodford N, Hawkey PM. 2008. Imported chicken meat as a potential source of quinolone-resistant *Escherichia coli* producing extended-spectrum beta-lactamases in the UK. J Antimicrob Chemother. 61:504–8.

29. Randall LP, Clouting C, Horton RA, Coldham NG, Wu G, Clifton-Hadley FA, Davies RH, Teale CJ. 2011. Prevalence of *Escherichia coli* carrying extended-spectrum β-lactamases (CTX-M and TEM-52) from broiler chickens and turkeys in Great Britain between 2006 and 2009. J Antimicrob Chemother. 66:86–95.

30. Radford AD, Noble PJ, Coyne KP, Gaskell RM, Jones PH, Bryan JG, Setzkorn C, Tierney Á, Dawson S. 2011. Antibacterial prescribing patterns in small animal veterinary practice identified via SAVSNET: the small animal veterinary surveillance network. Vet Rec. 169:310.

31. Wan NurIsmah WAK, Takebayashi Y, Findlay J, Heesom KJ, Jiménez-Castellanos JC, Zhang J, Graham L, Bowker K, Williams OM, MacGowan AP, Avison MB. 2018. Prediction of Fluoroquinolone Susceptibility Directly from Whole-Genome Sequence Data by Using Liquid Chromatography-Tandem Mass Spectrometry to Identify Mutant Genotypes. Antimicrob Agents Chemother. 62:e01814–17.

32. Tchesnokova V, Radey M, Chattopadhyay S, Larson L, Weaver JL, Kisiela D, Sokurenko EV. 2019. Pandemic fluoroquinolone resistant *Escherichia coli* clone ST1193 emerged via simultaneous homologous recombinations in 11 gene loci. ProcNatlAcadSci U S A. 116:14740–14748.

33. Findlay J, Gould VC, North P, Bowker KE, Williams MO, MacGowan AP, Avison MB. 2020. Characterization of cefotaxime-resistant urinary *Escherichia coli* from primary care in South-West England 2017-18. J Antimicrob Chemother. 75:65–71.

34. Mounsey O, Schubert H, Findlay J, Morley K, Puddy EF, Gould VC, North P, Bowker KE, Williams OM, Williams PB, Barrett DC, Cogan TA, Turner KM, MacGowan AP, Reyher KK, Avison MB. 2021. Infrequent Sharing of Fluoroquinolone-Resistant *Escherichia coli* from Dairy Farms as a Cause of Bacteriuria in Humans Living in the Same Geographical Region. BioRxiv

35. Pightling AW, Pettengill JB, Luo Y, Baugher JD, Rand H, Strain E. 2018. Interpreting Whole-Genome Sequence Analyses of Foodborne Bacteria for Regulatory Applications and Outbreak Investigations. Front Microbiol. 9:1482.

36. Ewers C, Bethe A, Semmler T, Guenther S, Wieler LH. 2012. Extended-spectrum β-lactamase-producing and AmpC-producing *Escherichia coli* from livestock and companion animals, and their putative impact on public health: a global perspective. Clin Microbiol Infect. 18:646–55.

37. Gandolfi-Decristophoris P, Petrini O, Ruggeri-Bernardi N, Schelling E. 2013. Extended-spectrum β-lactamase-producing Enterobacteriaceae in healthy companion animals living in nursing homes and in the community. Am J Infect Control. 41:831–5.

38. Timofte D, Maciuca IE, Williams NJ, Wattret A, Schmidt V. 2016. Veterinary Hospital Dissemination of CTX-M-15 Extended-Spectrum Beta-Lactamase-Producing *Escherichia coli* ST410 in the United Kingdom. Microb Drug Resist. 22:609–615.

39. Murray JK, Kinsman RH, Lord MS, Da Costa REP, Woodward JL, Owczarczak-Garstecka SC, Tasker S, Knowles TG, Casey RA. 2021. ‘Generation Pup’ – protocol for a longitudinal study of dog behaviour and health. BMC Vet Res. 17:1.

40. EUCAST. Breakpoint tables for interpretation of MICs and zone diameters. Version 8.1. http://www.eucast.org/fileadmin/src/media/PDFs/EUCAST_files/Breakpoint_tables/v_8.1_Breakpoint_Tables.pdf.

41. Bolger AM, Lohse M, Usadel B. 2014. Trimmomatic: a flexible trimmer for Illumina sequence data. Bioinformatics. 30:2114–2120.

42. Bankevich A, Nurk S, Antipov D, Gurevich AA, Dvorkin M, Kulikov AS, Lesin VM, Nikolenko SI, Pham S, Prjibelski AD, Pyshkin AV, Sirotkin AV, Vyahhi N, Tesler G, Alekseyev MA, Pevzner PA. 2012. SPAdes: a new genome assembly algorithm and its applications to single-cell sequencing. J Comput Biol. 19:455–477.

43. Seemann T. 2014. Prokka: rapid prokaryotic genome annotation. Bioinformatics. 30:2068–2069.

44. Zankari E, Hasman H, Cosentino S, Vestergaard M, Rasmussen S, Lund O, Aarestrup FM, Larsen MV. 2012. Identification of acquired antimicrobial resistance genes. J Antimicrob Chemother. 67:2640–2644.

45. Wirth T, Falush D, Lan R, Colles F, Mensa P, Wieler LH, Karch H, Reeves PR, Maiden MC, Ochman H, Achtman M. 2006. Sex and virulence in *Escherichia coli:*an evolutionary perspective. Mol Microbiol. 60:1136–1151.

46. Grüning B, Dale R, Sjödin A, Chapman BA, Rowe J, Tomkins-Tinch CH, Valieris R, Köster J; Bioconda Team. 2018. Bioconda: sustainable and comprehensive software distribution for the life sciences. Nat Methods. 15:475–476.

47. Connor TR, Loman NJ, Thompson S, Smith A, Southgate J, Poplawski R, Bull MJ, Richardson E, Ismail M, Thompson SE, Kitchen C, Guest M, Bakke M, Sheppard SK, Pallen MJ. 2016. CLIMB (the Cloud Infrastructure for Microbial Bioinformatics): an online resource for the medical microbiology community. Microb Genom. 2:e000086.

48. Stamatakis A, Ludwig T, Meier H. 2005. RAxML-III: a fast program for maximum likelihood-based inference of large phylogenetic trees. Bioinformatics. 21:456–463.

49. Stamatakis A. 2006. RAxML-VI-HPC: maximum likelihood-based phylogenetic analyses with thousands of taxa and mixed models. Bioinformatics. 22:2688–2690.

50. Argimón S, Abudahab K, Goater RJE, Fedosejev A, Bhai J, Glasner C, Feil EJ, Holden MTG, Yeats CA, Grundmann H, Spratt BG, Aanensen DM. 2016. Microreact: visualizing and sharing data for genomic epidemiology and phylogeography. MicrobGenom. 2:e000093.

